# Targeting PPT1 with ezurpimtrostat sensitives liver tumor to immunotherapy by switching cold into hot microenvironments

**DOI:** 10.1101/2023.01.18.524541

**Authors:** Eloïne Bestion, Madani Rachid, Annemilaï Tijeras-Raballand, Gael Roth, Thomas Decaens, Christelle Ansaldi, Soraya Mezouar, Eric Raymond, Philippe Halfon

## Abstract

**Background:** Palmitoyl-protein thioesterase-1 (PPT1) is an exciting druggable target for inhibiting autophagy in cancer.

**Methods:** In this study, we aimed to evaluate the effects of ezurpimtrostat-targeting PPT1 in combination with an anti-PD-1 antibody in liver cancer using a transgenic immunocompetent mouse model.

**Results:** Herein, we revealed that inhibition of PPT1 using ezurpimtrostat, a safe anticancer drug in humans, decreased the liver tumor burden by inducing the penetration of lymphocytes within tumors when combined with anti-programmed death-1 (PD-1). Inhibition of PPT1 potentiates the effects of anti-PD-1 immunotherapy by increasing the expression of major histocompatibility complex (MHC)-I at the surface of liver cancer cells and modulates immunity through recolonization and activation of cytotoxic CD8^+^ lymphocytes.

**Conclusions:** Ezurpimtrostat turns cold into hot tumors and, thus, constitutes a powerful strategy to improve T cell-mediated immunotherapies in liver cancer.

**Summary box:** We reported that inhibiting palmitoyl-protein thioesterase-1 enzyme (PPT1) enhances the antitumor activity of anti-programmed death-1 (PD-1) in liver cancer in preclinical models. This study provides the rational for this combination in cancer clinical trials.

**Graphical abstract:** 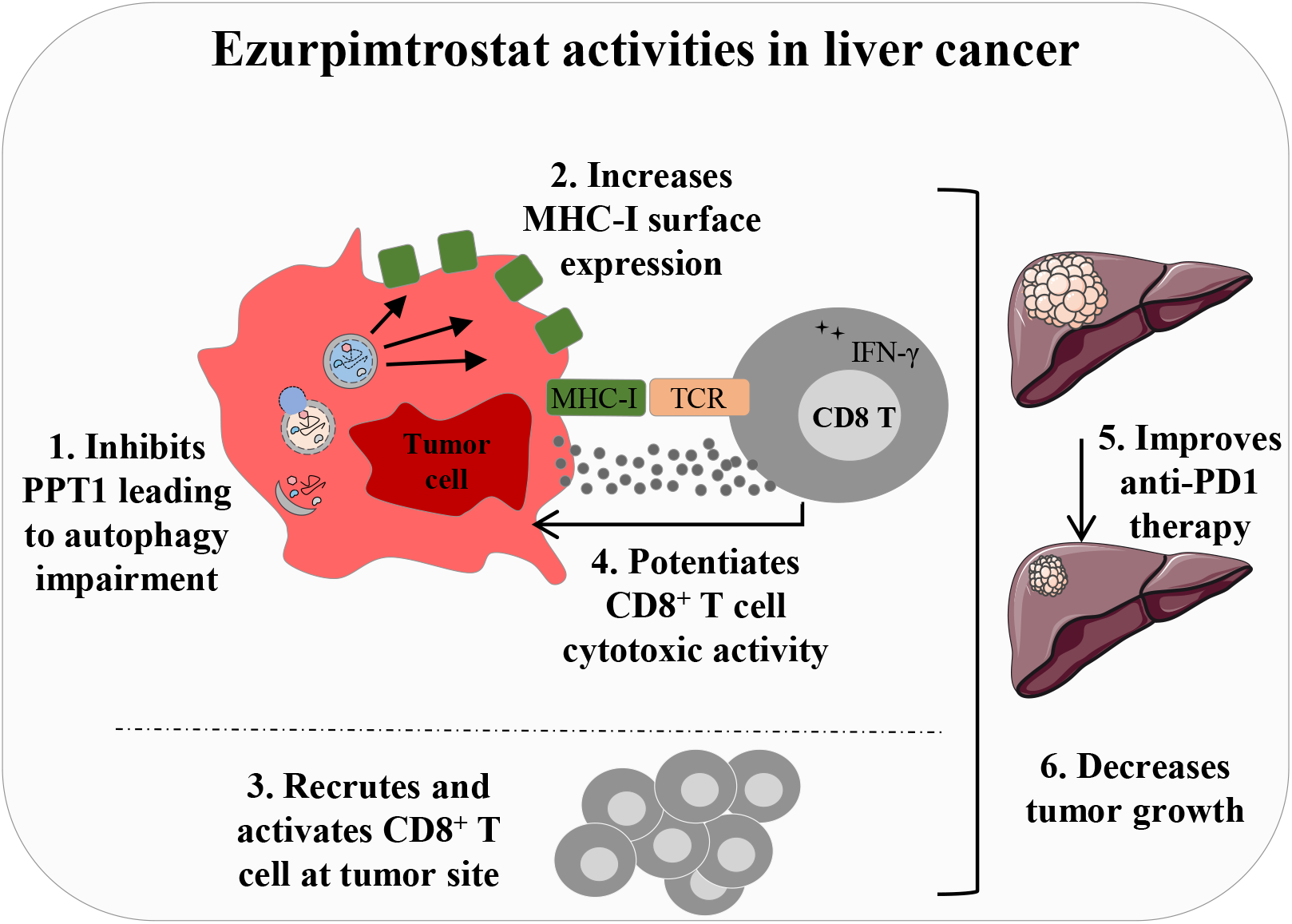

**Ezurpimtrostat activities in cancer:** The absence of immune effectors especially cytotoxic cells in the microenvironment of cold tumor is associated with a lack of response to ICI. This condition is mainly due to an increase in the autophagy process responsible for the sequestration and destruction of an antigen-presenting molecule, MHC-I. The inhibition of PPT1 using ezurpimtrostat treatment led to (**1**) the inhibition of PPT1 and consequently the autophagy process, (**2**) the increase of MHC-I surface expression, and (**3**) the recruitment and the activation of CD8^+^ T cells at tumor site leading to (**4**) the improvement of CD8^+^ T cell cytotoxic activity. Thus, ezurpimtrostat-treated tumors become eligible for anti-PD-1 immunotherapy as the combination of both led to decreased macronodules, micronodules, and tumor growth.

## Background

Palmitoyl-protein thioesterase 1 (PPT1), a glycoprotein belonging to the palmitoyl-protein thioesterase family, has been shown to play a central role in the control of cellular autophagy and has been reported to be highly expressed in several cancer cell lines as well as in tissues from patients with advanced cancers associated with poor survival (1,2). PPT1 has been investigated as a potential target for cancer therapies (3). We and others have reported that PPT1 inhibition can limit tumor growth in several cancers (1,2,4). Recently, we highlighted that PPT1 inhibition leads to lysosomal dysregulation, late-stage autophagy inhibition, and cancer cell death in hepatocarcinoma (HCC) models (4). In humans, targeting PPT1 is safe and associated with signal of antitumor activity in HCC (5).

Cancer management has been revolutionized using a therapeutic class of monoclonal antibodies and immune checkpoint inhibitors (ICI). Among them, the anti-programmed death-1 (PD-1) immune checkpoint blockade has been used to enhance the effector function of tumor-specific CD8^+^ T cells (6). In an immune-rich environment, HCC is a major indication for immune-based therapies. However, despite being associated with response rates in the range of 15 to 20% in phase 3 studies in first- and second-line settings, single-agent ICI did not significantly improve overall survival (7–11). Moreover, non-responders progress at a rate similar to that observed for the natural history of the disease, and longer follow-up of clinical populations revealed that late relapses emerge, suggesting acquired resistance to ICI monotherapy (12). Combination therapy using ICI in combination with other anticancer agents modulates the tumor microenvironment in preclinical data, providing a rationale for developing novel anticancer strategies for advanced unresectable HCC. For example, treatment with atezolizumab plus bevacizumab was associated with significantly better overall and progression-free survival outcomes than sorafenib in patients with HCC not previously treated with systemic therapy and has become the new standard of care in patients with HCC (9). Interestingly, the inhibition of PPT1 was found to increase antigen presentation and was associated with CD8^+^ T cell tumor colonization, proliferation, and activation involved in tumor cell killing (13) enhancing the antitumor efficacy of ICI in melanoma cancer (14).

Ezurpimtrostat/GNS561, a novel chemical entity, is a lysosomotropic small basic lipophilic molecule and a potent PPT1 inhibitor. Several findings highlight the interest in ezurpimtrostat treatment in cancer: (1) blocking PPT1-dependent autophagic activity; (2) inducing lysosomal deacidification, unbound zinc accumulation, and permeabilization of lysosomal membranes; (3) activating caspases and cancer cell lethality; and (4) preliminary signal of activity at the clinical stage with a favorable safety profile (4,5,15). In this study, we aimed to evaluate the effects of ezurpimtrostat-targeting PPT1 in combination with an anti-PD-1 antibody in liver cancer using a transgenic immunocompetent mouse model. Our study reported that inhibition of PPT1 using ezurpimtrostat decreased tumor growth when combined with PD-1. Ezurpimtrostat potentiates ICI by increasing the expression of major histocompatibility complex (MHC)-I protein at the surface of cancer cells and modulating immunity by recolonization and activation of cytotoxic CD8^+^ lymphocytes. This study provides evidence that the inhibition of PPT1 sensitizes tumors to immunotherapy by switching cold into hot tumors, highlighting the clinical rationale for combining PPT1 inhibitors and ICI in cancer.

## Methods

### Mice model and treatment

Seven-week-old transgenic C57BL6/ASV-B male mice obtained from TAAM UAR44 (CNRS, Orléans) developing HCC were described previously (40). Briefly, precise targeting of the SV40 T early region expression in the liver of transgenic mice was achieved using 700 base pairs of antithrombin regulatory sequences to control oncogene expression. Hepatocyte hyperplasia/dysplasia occurred at 8 weeks (W), nodules corresponding to the adenomatous stage were observed in W12, and diffuse HCC from W16. HCC progression is also characterized by angiogenesis and severe vessel anomalies, such as capillarization and arterialization. All experiments were performed following Directive 2010/63/EU of the European Parliament and Council on September 22, 2010. This project was approved by the local ethic committee (Comité d’éthique en experimentation animale Lariboisière-Villemin n°9).

Before the experiment, a control ultrasound procedure was performed to allocate animals homogeneously based on the liver volume. As illustrated in **Fig. 1**, the study design included the following 10 animals: vehicle (control group), ezurpimtrostat, anti-PD-1 (CVP033, CrownBio), and combination group (ezurpimtrostat with anti-PD-1). For ezurpimtrostat (alone or in combination, 50 mg/kg) and vehicle, the mice were treated daily, 6 days/week, at a constant dosage of 10 ml/kg *per os*. Anti-PD-1 was diluted in phosphate-buffered saline (PBS; Life Technologies, L0615-500) extemporaneously, before administration. Mice receiving anti-PD-1 (alone or in combination, 10 mg/kg) were treated twice a week, using a constant dosage of 10 ml/kg *via* an intraperitoneal route. All mice were weighed three times per week to adjust the volume of the treatment.

**Fig. 1.**
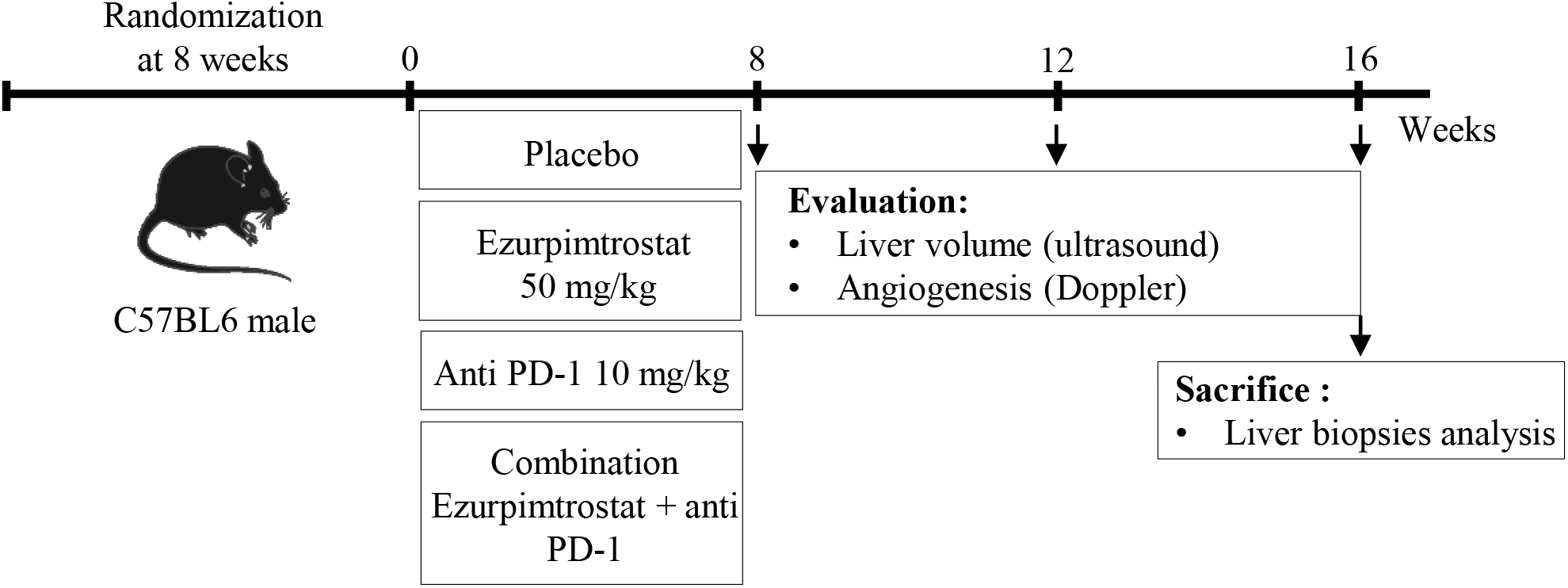
Study design. Four groups of mice (n=10) were treated for 8 weeks with ezurpimtrostat (10 mg/kg/6 × week, *per os*), anti-PD-1 (CVP033, 10 mg/kg/2 × week intraperitoneal), ezurpimtrostat (10 mg/kg/6 × week) plus anti-PD-1 (CVP033, 10 mg/kg/2 × week intraperitoneal), or vehicle (water at pH 4 ± 0.2, *per os*) as a control. Every 4 weeks, echography and Doppler were performed to monitor tumor burden. At week 16, the animals were sacrificed, and organs were harvested for preservation.

Mice were euthanized at W16, and livers were collected, weighed, macronodule-numbered, and then paraffin-embedded for analysis. All treatments were well tolerated based on animal behavior and body weight follow-up (**Supplementary Fig. S1**). One animal was excluded from the analyses because the animal died prematurely in the ezurpimtrostat group due to the mishandling gavage procedure.

### Ultrasound and Doppler

Ultrasound Doppler assessment in this model has been described previously (41). In this model, liver tumor angiogenesis has been described to be correlated with blood flow in the celiac trunk (CT). When abnormal angiogenesis is sprayed into the liver, blood flow increases in CT. Ultrasound imaging was performed every 4 weeks (**Fig. 1**). The minimum detectable velocity was approximately 1 cm/s and the maximum was 120 cm/s.

### Immunostaining

Immunohistochemical staining was performed on 5-μm-thick slices from paraffin-embedded tumors and counterstained with hematoxylin-phloxin-saffron (HES), using an automated immunohistochemical stainer (Tissue-Tek Prisma^®^ Plus, Sakura, France) following the manufacturer’s instructions. Immunostaining was performed using an automated immunohistochemical stainer (BondMax; Leica, France). CD8 antibody (Abcam, #ab209775, 1/200) was used to assess lymphocyte localization in HCC livers at perinodular and intranodular levels.

The immunofluorescence procedure was performed on 5-μm-thick slices from paraffin-embedded tumors. Tissue sections were deparaffinized in xylene baths 3 times for 5 min each before gradual hydration with ethanol in successive 5-min baths of 2 min in 100%, 95%, 70%, and 50% ethanol. The tissue sections were rinsed twice for 5 min in phosphate-buffered saline (PBS). p62 and PPT1 labeling was preceded by a heat-induced epitope retrieval step to improve the detection of antibody staining in paraffin-embedded tissue sections. Coverslips were placed in boiling antigen retrieval buffer (Tris 10 mM, EDTA 1 mM, Tween 20 0.05% in water, pH 9) for 10 min and then immersed in cold water to allow the antigenic sites to re-form. The coverslips were then washed once in PBS before two permeabilization baths with permeation buffer (Triton 0.4% in PBS) for 10 min at room temperature (RT). The coverslips were washed twice with PBS. For all antibodies, specific sites were blocked with a blocking buffer solution (5% fetal bovine serum [FBS, Life Technologies, #10270-098], 0.1% Tween 20 [Biosolve, #20452335] in PBS [Life Technologies, #14200-067]) for 30 min at RT and washed twice. p62 (Abcam, #ab240635, 1/150), MHC-I (BioLegend, #125508, 1/150), PPT1 (Invitrogen, #PA5-29177, 1/220) antibodies, and 4’,6-diamidino-2-phenylindole (DAPI, Invitrogen, #D1306, 1/1000) were incubated overnight at 4°C in the dark. The following day, coverslips were washed twice with PBS and incubated for 1 h at RT with Alexa647-conjugated anti-rabbit secondary antibody (Invitrogen, #A-31573, 1/500) to reveal both p62 and PPT1 proteins. All antibodies were diluted in a blocking buffer solution. Labeled cells were washed twice with PBS, mounted using Fluoromount™Aqueous Mounting Medium, and stored overnight at 4°C before analysis. For image analysis, fluorescence images were acquired using an ApoTome module associated with a Zeiss microscope (Zeiss, Germany) equipped with an AxioCam MRm camera and collected using the AxioVision software (Zeiss, Germany) with a 63× oil objective. Randomly selected microscopic tiles (5 × 5) were acquired using the Zen 3.0 software (Blue Edition, Zeiss, Germany). Labeled areas from selected fields were quantified using ImageJ software.

### Cell isolation and cell line culture

The HepG2 (hepatocellular carcinoma, ATCC, #HB-8065) cell line was cultured in DMEM low glucose (Thermo Fisher Scientific, #21,885,025) supplemented with 1% penicillin-streptomycin (Dutscher) and 10% FBS (GE Healthcare, #SV30160.03C).

Peripheral blood mononuclear cells (PBMCs) were isolated from buffy coats of healthy donors by density-gradient centrifugation using Ficoll (Eurobio, #CMSMSL01-01) at 800 x g (brake off) for 30 minutes at 20°C. PBMCs were cryopreserved at −80°C in 90% FBS (Gibco, #10099141) and 10% dimethyl sulfoxide (DMSO, Sigma-Aldrich, #D2650). Lymphocytes were isolated and purified using a Pan T Cell Isolation Kit (Miltenyi Biotech, #130-096-535). The degree of lymphocyte purity achieved was at least 98% and was assessed using standard sorting procedures by flow cytometry.

### Flow cytometry analysis

IFN-γ expression was evaluated in the CD3^+^CD8^+^ lymphocyte population co-cultured with HepG2 cells and treated or not with ezurpimtrostat at 0.6, 6, or 7 μM for 24 h. After incubation, the cells were suspended in PBS (Dutscher, #L0615-500) containing 1% FBS and 2mM EDTA (Dutscher, #P10-15100). Cells were labeled with antibodies against viability fixable dye (Miltenyi Biotec, #130-109-812), CD3 (Miltenyi Biotec, #130-113-141), CD8 (Miltenyi Biotec, #130-110-681), and IFN-γ (BD Pharmingen, #560741).

MHC-1 expression in HepG2 cells was evaluated. Briefly, cells (60,000 cells/well) were plated in a 96-well tissue culture plate in 90 μL DMEM with low glucose. Twenty-four hours after plating, the cultivated cells were treated with increasing concentrations of GNS561 (10 μL) or vehicle control and were then incubated for an additional 24 h. The cells were harvested, washed, and incubated with v450 viability stain (Invitrogen, #650863-14), MHC-I (Invitrogen, #65-0863), and NBR1 (Santa Cruz Biotechnology, #sc130380) antibodies.

Labeled cells were acquired from at least 50,000 cells, collected using an Attune NxT flow cytometer (Thermo Fisher), and analyzed using FlowJo software (FlowJo v10.6.2).

### Statistical analysis

Statistical analyses were performed using the Prism 8.4.3 software. For datasets with a normal distribution, multiple comparisons were performed using one-way ANOVA with Tukey’s post-hoc analysis. An unpaired t-test was used to compare two groups of data with normal distributions. Data are presented as mean ± standard deviation (SD). Statistical significance was defined as a p-value of < 0.05.

## Results

### PPT1 inhibition in combination with anti-PD-1 decreases tumor growth

As autophagy inhibitors in combination with ICI provide opportunities for enhancing antitumor activity (13,14), we investigated the efficacy of ezurpimtrostat in combination with anti-PD-1 in a transgenic immunocompetent HCC mouse model according to the study design described in **Fig. 1**. Twelve weeks post-treatment, ezurpimtrostat monotherapy or in combination with ICI led to a significant decrease in liver volume compared to the vehicle control or anti-PD-1 group (**Table 1**). Moreover, the tumor volume at the end of the experiment (day 16) was significantly smaller for mice treated with ezurpimtrostat monotherapy or in combination with ICI than for the vehicle control or anti-PD-1 group. At both time points, no differences were observed between the control and anti-PD-1 treatment groups. As HCC has been reported to be a highly vascular tumor (16), we next evaluated blood flow at 12 and 16 weeks. As illustrated in **Table 1**, ezurpimtrostat monotherapy or in combination with ICI significantly decreased blood flow compared to the vehicle control, while an increase was found during anti-PD-1 treatment. Interestingly, at the end of the experiment, only the combination of ezurpimtrostat and anti-PD-1 led to a significant decrease in blood flow in the liver compared with the vehicle control. Taken together, these findings suggest that the antitumor activity of ezurpimtrostat was enhanced by its combination with anti-PD-1.

**Table 1.**
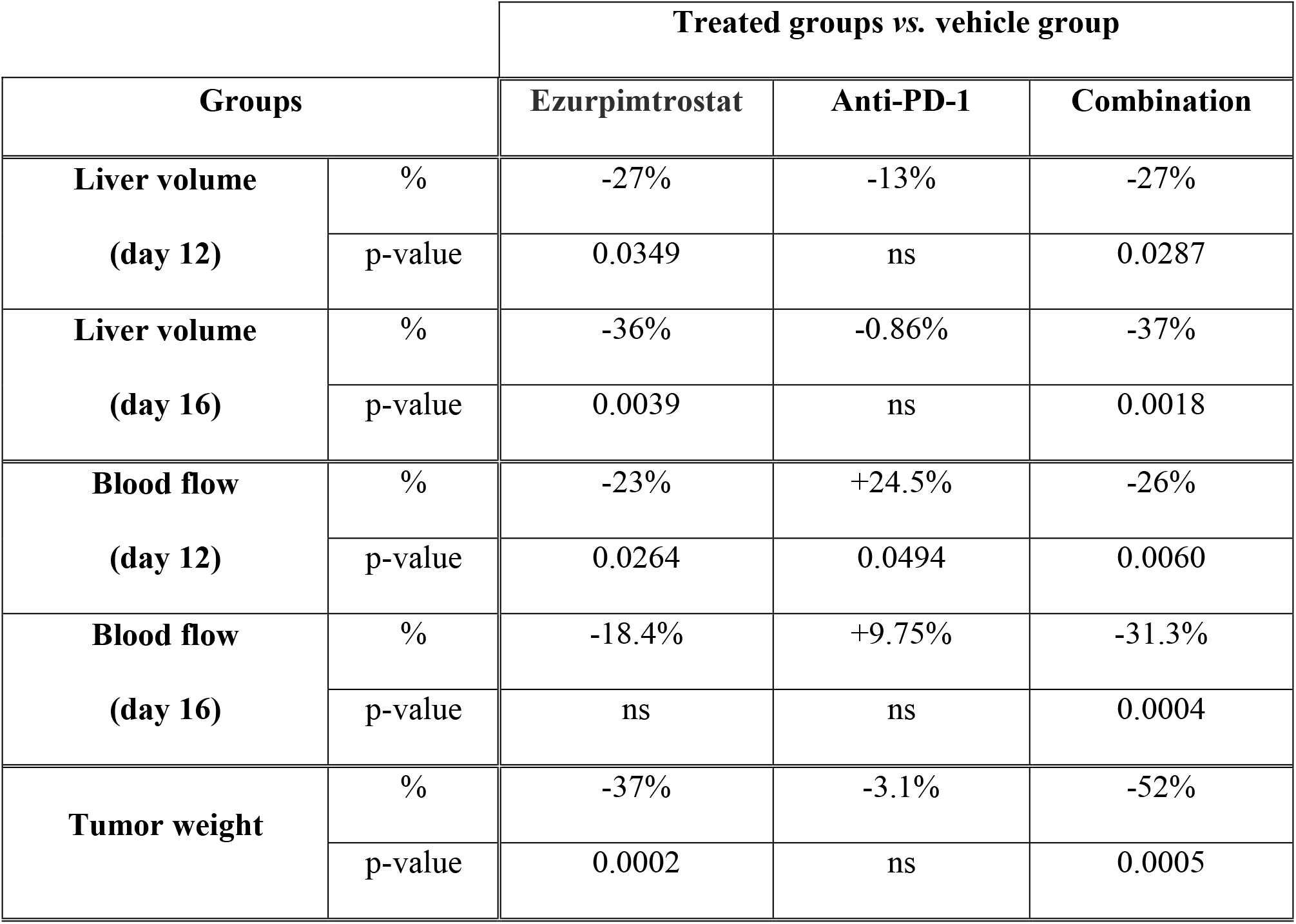
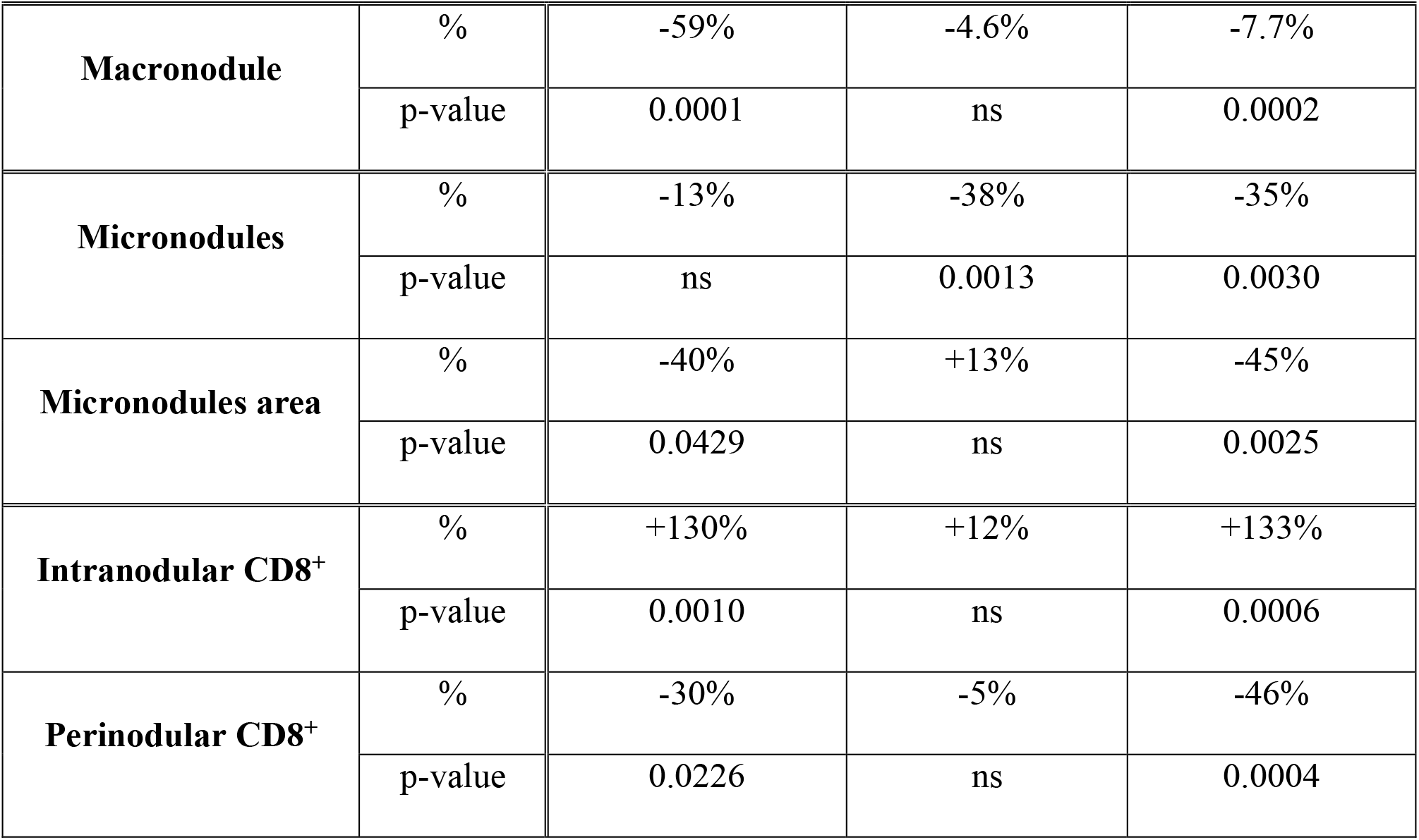
Parameters evaluated in mice treated or not by ezurpimtrostat, anti-PD-1, or the combination of compounds. ns: not significant

### PPT1 inhibitor combined with anti-PD-1 affects neoplastic nodules

As illustrated in **Fig. 2A**, the livers of mice treated with ezurpimtrostat monotherapy or anti-PD-1 plus ezurpimtrostat combination were similar in appearance, with a smaller liver size and fewer macronodules than those in the control and anti-PD-1 groups. Indeed, ezurpimtrostat treatment significantly decreased liver weight and macronodule number when administered as monotherapy or in combination (**Fig. 2B** and **Fig. 2C**; **Table 1**), whereas no significant difference was observed between the vehicle and anti-PD-1 groups. Interestingly, the number of macronodules significantly decreased when mice were treated with ezurpimtrostat in combination with anti-PD-1 compared to ezurpimtrostat alone, suggesting a benefit of the combination (**Fig. 2C**). Next, the variation in micronodule number and area was investigated using hematoxylin-phloxin-saffron coloration in the excised liver (**Fig. 2D**). The combination treatment showed greater efficiency in reducing both the number (**Fig. 2E**) and area (**Fig. 2F**) of the neoplastic micronodules compared to the control (**Table 1**), suggesting that inhibition of PPT1 potentiates the effects of anti-PD-1 immunotherapy.

**Fig. 2.**
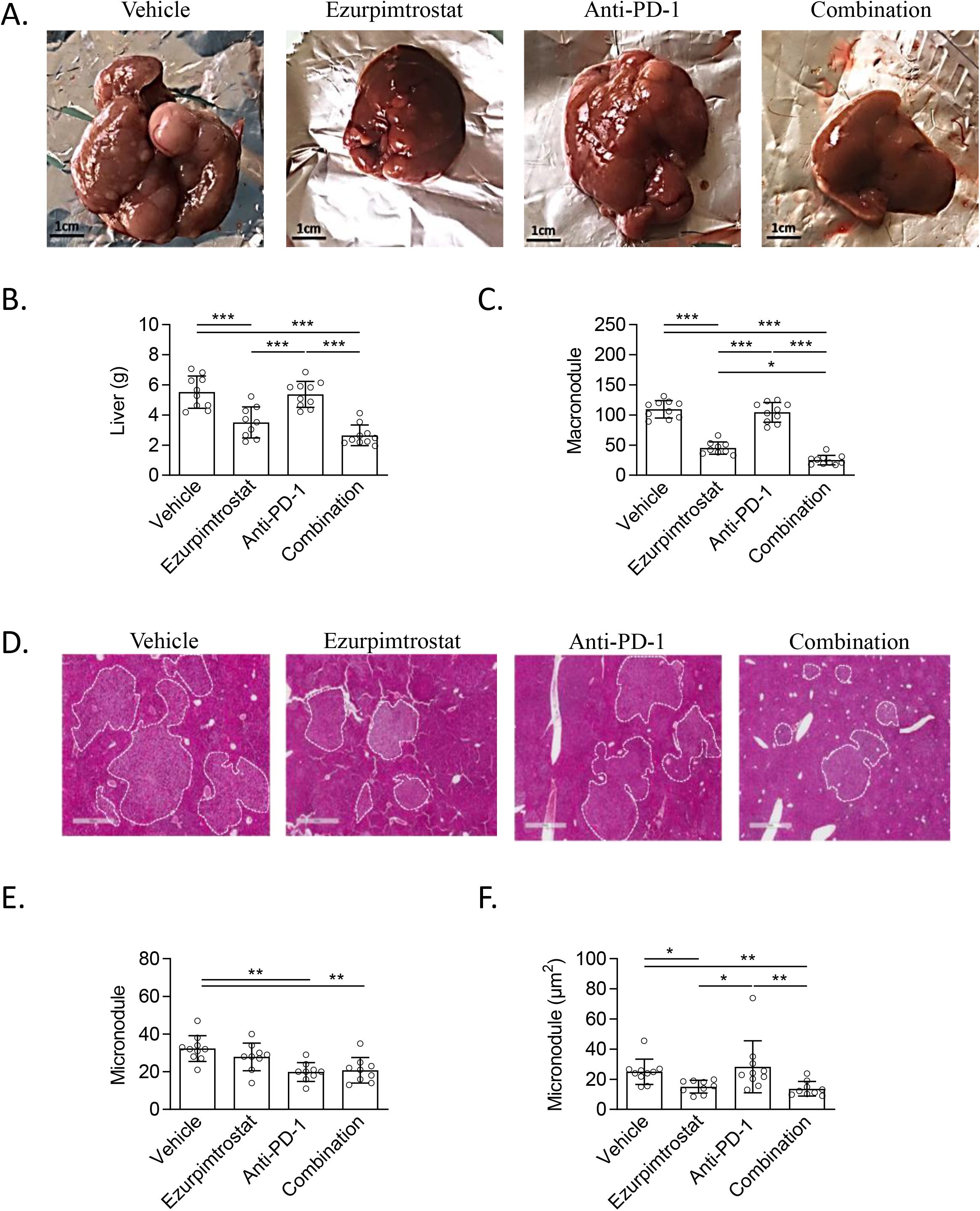
Ezurpimtrostat in combination with anti-PD-1 affects neoplastic nodules. (**A**) Representative pictures illustrating macroscopic livers from each group. Graph showing (**B**) liver weight and (**C**) macronodule number at the surface of the liver for each group and expressed as the mean ± standard deviation of 10 fields. *\*p<0.05, ***p<0.001*. (D) Images illustrating micronodules from the livers of each group. (**E**) Number and (**F**) area of micronodules were quantified on excised livers and expressed as the mean ± standard deviation. *\*p<0.05, **p<0.01*.

### Local autophagy inhibition facilitates T cell tumor colonization and activation

To evaluate the local action of ezurpimtrostat at the tumor level, we first evaluated two key mechanisms related to ezurpimtrostat action: inhibition of autophagy through the expression of p62 and inhibition of PPT1 at the tumor level. As illustrated in **Fig. 3**, we observed an increase in p62 (*p=0.0286;* **Fig. 3A**) and a decrease in PPT1 (*p=0.0153;* **Fig. 3B**) expression, highlighting the local action of ezurpimtrostat in inhibiting autophagy and PPT1.

**Fig. 3.**
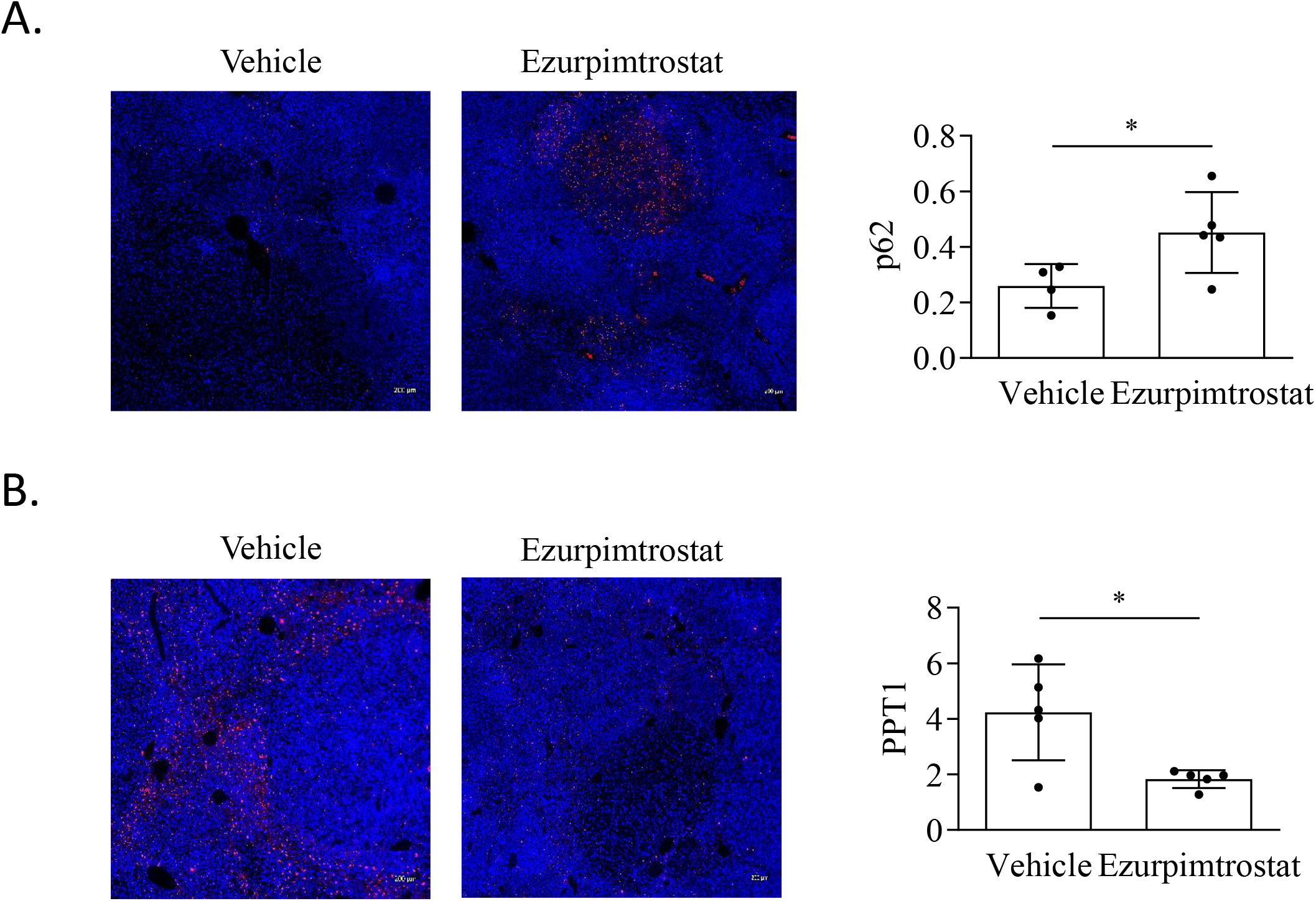
Ezurpimtrostat inhibits autophagy and PPT1 expression at the tumor level. (**A**) Representative pictures of p62 (red) expression and nucleus (blue) in liver biopsies from mouse treated by vehicle (left panel) as control or ezurpimtrostat (right panel); and ratio of p62 expression to cell nucleus of slides from 5 mice for each groups using area analysis (ImageJ). Data represent the mean ± standard deviation. *\*p<0.05*. (**B**) Representative pictures of PPT1 (red) expression and nucleus (blue) in liver biopsies from a mouse treated by vehicle (left panel) as control or ezurpimtrostat (right panel); and ratio of PPT1 expression to cell nucleus of slides from 5 mice for each groups using RawIntDen (ImageJ) as a measure of fluorescence intensity. Data represent the mean ± standard deviation. *\*p<0.05*.

Next, we investigated the effects of ezurpimtrostat on the tumor cell microenvironment. As illustrated in **Fig. 4A**, we observed the presence of CD8^+^ T cell at intra- and perinodular levels. Ezurpimtrostat treatment alone or in combination led to a significant increase in the CD8^+^ T cell population at the intranodular level, while the same population was significantly under-represented at the perinodular level (**Fig. 4B** and **Fig. 4C**; **Table 1**). In contrast, anti-PD-1 treatment alone did not affect lymphocytes’ recruitment, either at the intra- or perinodular levels. To investigate the effect of ezurpimtrostat on lymphocytes, we performed a co-culture experiment. Interestingly, *in vitro* ezurpimtrostat treatment of the HepG2 HCC cell line co-cultured with different ratios of lymphocytes led to a decrease in cancer cell viability (**Fig. 4D**), suggesting that ezurpimtrostat acted on lymphocytes. To assess the immunomodulatory role of ezurpimtrostat on lymphocytes, we investigated CD8^+^ T-cell activation in co-culture experiments. Indeed, ezurpimtrostat led to a dose-dependent increase of interferon (IFN)-γ expression by lymphocytes co-cultured with HepG2, indicating that the compound activated immune cells (**Fig. 4E**).

**Fig. 4.**
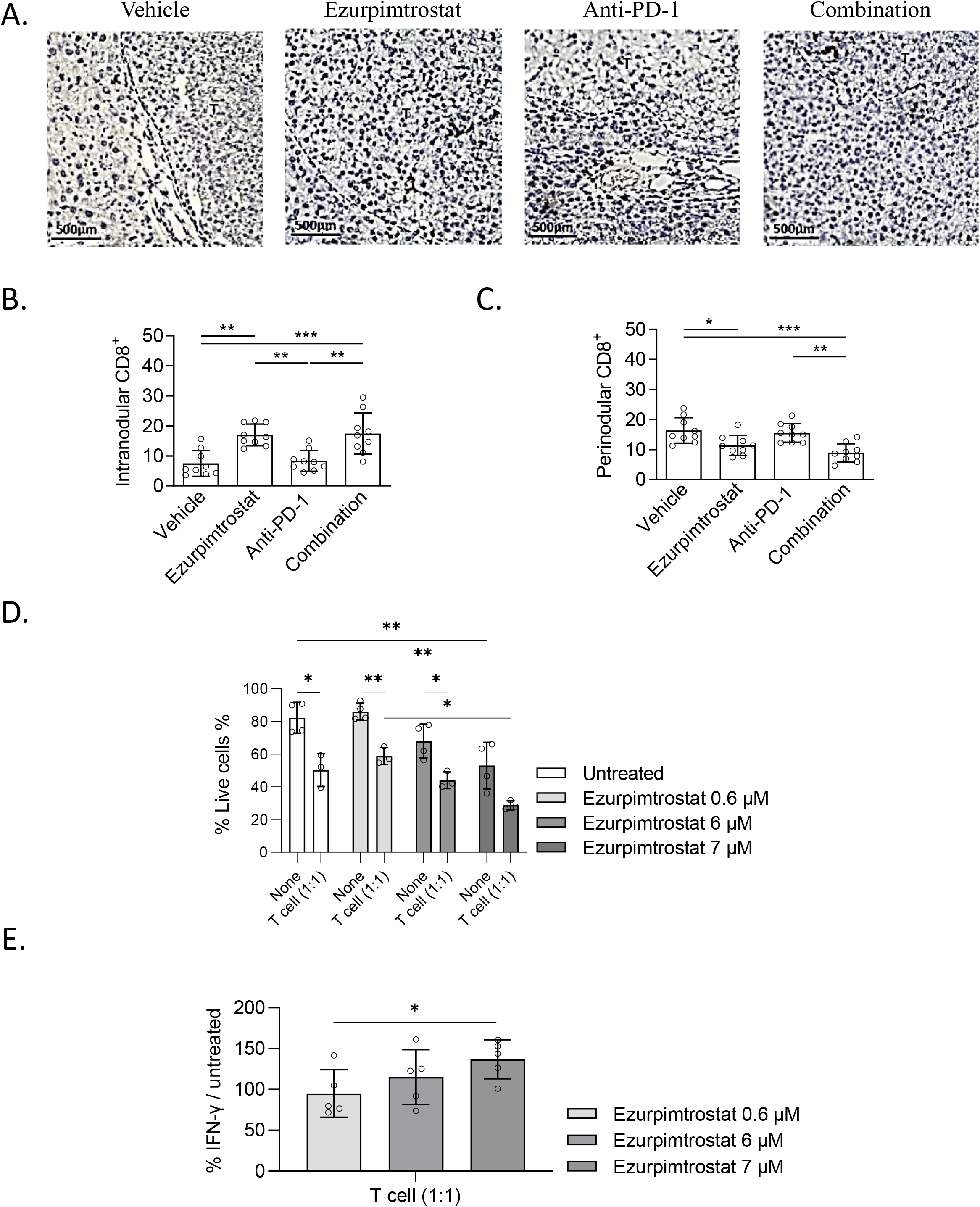
Ezurpimtrostat led to tumor MHC-I expression and immune T cell tumor colonization. (**A**) Representative pictures illustrating CD8^+^ T cell population from excised liver. Quantification of CD8^+^ T cell population at the (**B**) intranodular and (**C**) perinodular localization. Each spot represented the mean ± standard deviation of 10 fields. *\*p<0.05, **p<0.01, ***p<0.001*. (**D**) Mean fluorescence intensity of IFN-γ expressed by CD3^+^CD8^+^ lymphocytes co-cultured with cancer cell line at 0.5, 1, or 2 ratio and treated or not for 24 h by ezurpimtrostat at 0.6, 6, or 7 μM. Data represent the mean ± standard deviation. (**E**) Cell viability of cancer cells co-cultured or not with lymphocytes at 0.5, 1, or 2 ratio and treated or not for 24 h by GNS561 at 0.6, 6, or 7 μM. Data represent the mean ± standard deviation. *\*p<0.05*.

Taken together, these findings indicate that autophagy inhibition through ezurpimtrostat treatment is crucial in (**1**) tumor recolonization of cytotoxic CD8^+^ lymphocytes and (**2**) their activation to potentialize (**3**) the antitumor activity of the compound.

### Local autophagy inhibition restores tumor cell MHC-I expression

CD8^+^ effector T cells exert antigen-specific cytotoxic effects on tumor cells by recognizing tumor antigens carried by MHC-I class I molecules (17). Here, we report a significant increase in MHC-I protein expression in tumors from mice treated with ezurpimtrostat compared to the control (*p=0.0231;* **Fig. 5A**). Moreover, ezurpimtrostat treatment of cancer mice resulted in a positive correlation between the expression of p62, reflecting the inhibition of autophagy, and MHC-I at the tumor level (**Fig. 5B**). No correlation was observed in the untreated control group, suggesting that the inhibitory activity of ezurpimtrostat on the autophagy pathway was directly related to the modulation of tumor-level MHC-I protein expression. More specifically, MHC-I protein expression was found to be increased *in vitro* in HepG2 cells treated with ezurpimtrostat in a dose-dependent manner, *in vitro* (**Fig. 5C**), highlighting the specificity of the compound in modulating MHC-I protein expression in cancer cells. In contrast, ezurpimtrostat significantly decreased in a dose-dependent manner the expression of the neighbor of *BRCA1* gene 1 (NBR1) (**Fig. 5D**), which was previously identified as an autophagy cargo receptor involved in the sequestration and destruction of MHC-I protein (13).

**Fig. 5.**
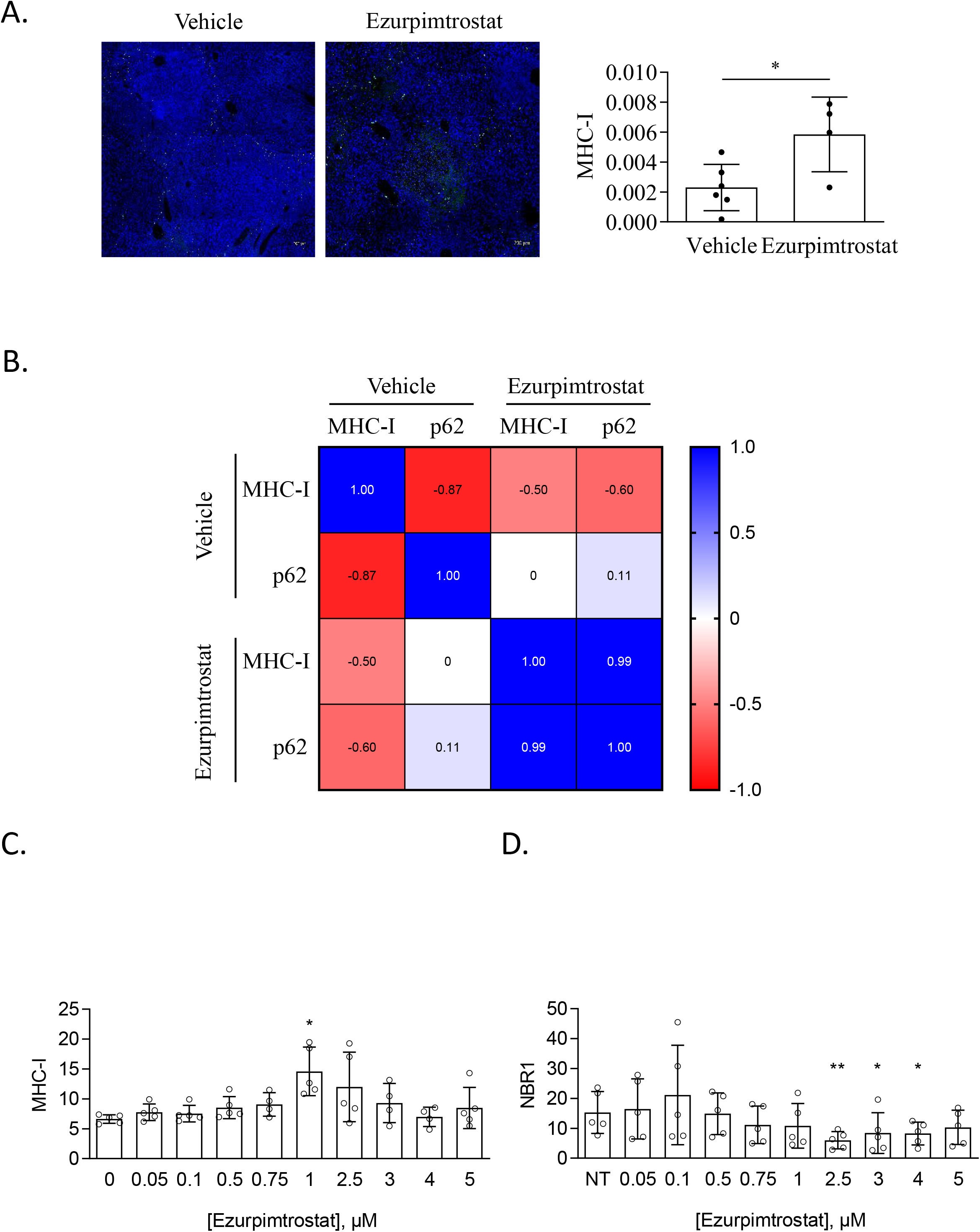
Ezurpimtrostat restores MHC-I expression on cancer cells. (**A**) Representative pictures (left panel) and graph (right panel) of MHC-I expression (green) and nucleus (blue) in liver biopsies from a mouse treated by vehicle (left panel) as control or GNS561 (right panel). Using RawIntDen (ImageJ) as a measure of fluorescence intensity, level of expression was expressed as the ratio of MHC-I expression to cell nucleus of slides from 4 mice for each group. Data represent the mean ± standard deviation. *\*p<0.05*. (**B**) Correlation matrix between MHC-I and p62 expression from the control and GNS561 groups. Scatter dot plot illustrating the mean fluorescence intensity of (**C**) MHC-I and (**D**) NBR1 on HepG2 cell line treated or not by increase concentration of GNS561 during 24 h. Data represent the mean ± standard deviation. *\*p<0.05, **p<0.01*.

## Discussion/Conclusion

This study reports that PPT1 inhibition using ezurpimtrostat in combination with anti-PD-1 impairs tumor growth and angiogenesis, significantly reducing the number of macronodules in liver cancer, when anti-PD-1 alone does not present a significant antitumor effect (**Fig. 6**). The inhibition of PPT1 and mechanisms related to autophagy potentiate the effects of anti-PD-1 immunotherapy by increasing tumor MHC-I expression, allowing the recovery of antigen recognition and recruitment of cytotoxic CD8^+^ lymphocytes. Although autophagy acts as a suppressor of early tumorigenesis (18), it also contributes to tumor initiation and growth, as well as cancer cell development and proliferation in various tumor types (18). Autophagy is involved in immune evasion (19,20) and promotes resistance to anti-PD-1-based therapeutic strategies/immunotherapy (21). Our data showed that PPT1 inhibition by ezurpimtrostat sensitizes tumors to ICI.

Ezurpimtrostat combined with anti-PD-1 led to T-cell recolonization at the tumor site, suggesting that PPT1 inhibition counteracted immune evasion. In contrast, the combination of anti-PD-1 and PPT1 inhibition by hydroxychloroquine did not modulate the number of T cells (14) suggesting that ezurpimtrostat has potent immunomodulatory activity in cancer. In addition, PPT1 inhibition modulates infiltrating-CD8^+^ T cells activation through the expression of IFN-γ, a key cytotoxic cytokine involved in tumor cells’ apoptosis (22,23). This finding is in accordance with the increased production of IFN-γ by tumor-infiltrating CD8^+^ cells observed in B16 melanoma cells and H22 tumor ascites models (24). Interestingly, the authors reported that T-cell production of IFN-γ during chloroquine treatment was mediated by tumor-associated macrophages (TAMs). TAMs have been extensively investigated in cancer, especially their polarization in the M1 (pro-inflammatory) and M2 (anti-inflammatory) phenotypes (25) which exhibit anti-tumoral and pro-tumoral activities, respectively (26). Sharma *et al*. reported that inhibition of autophagy targeting PPT1 leads to a switch in macrophage polarization from M2 to M1 (14). The key role of macrophage polarization switching after PPT1 inhibitors has been previously reported (14,24). Although macrophages have been described as key immune cells involved in tumor suppression, data on their role in tumor immune microenvironment modulation are increasing in the literature. Indeed, the switch from M2 to M1 polarization allows IFN-β secretion by pro-inflammatory macrophages (14) as well as the tumor colonization of myeloid-derived suppressor cells and Treg cells (24), contributing to T-cell cell-mediated cytotoxicity. However, the level of synergistic action of ICI on the modulation of these immune cells remains to be determined.

The results of this study are consistent with previous reports by Yamamoto *et al*., showing that the activation of CD8^+^ T lymphocytes strongly depends on interactions with MHC-I at the surface of tumor cells to efficiently recognize antigens (13). However, ezurpimtrostat specifically modulated MHC-I protein expression in cancer cells dose-dependently *in vitro* and *in vivo*. The regulatory mechanism of MHC-I and its upregulation were previously involved in the restriction of immune escape (27) and its reduction was associated with a decrease in T cell-mediated killing of cancer cells (28). PPT1 inhibition in pancreatic cancer models overexpresses MHC-I proteins that potentiate cancer immunotherapy (13). In contrast, Zeng *et al*. reported that MHC-I protein expression decreased after PPT1 inhibition in non-small cell lung cancer cells treated by radiation therapy (29). Moreover, the authors highlighted that MHC-I protein expression is positively correlated with the recruitment of CD8^+^ T cells. Interestingly, this study showed that ezurpimtrostat treatment led to the tumor colonization of CD8^+^ T cells. Other mechanisms may be involved, such as the key role of antigen-presenting cells, such as dendritic cells, which express more MHC-I proteins at their surface after anti-autophagy drug treatment. This was, for instance, described as promoting the cytotoxic response of CD8^+^ T cells during the viral immune response (30). MHC-I is a key biomarker in cancer, as the decrease in its expression has been correlated with a worse prognosis (31), and mutation or loss of its heterozygosity has been implicated in resistance to ICI therapy (32–34).

Ezurpimtrostat treatment led to decreased NBR1 protein expression in cancer cells. In addition to the NBR1 mRNA expression found in most cancers, it was associated with low cancer specificity (35,36). NBR1 overexpression increased the proliferation of renal cancer cells *in vitro* (37) and promoted tumor metastasis in a breast cancer mouse model (38). In contrast, the knockdown of NBR1 inhibits *in vitro* cell migration (39) suggesting that NBR1 is involved in cancer development. Here, we report that ezurpimtrostat treatment led to a decrease in NBR1 protein levels, which is consistent with previous reports by Yamamoto *et al*., highlighting the key role of NBR1 in immune evasion. Future research must evaluate NBR1 as a potential target for therapeutic strategies.

To overcome primary and acquired resistance, several clinical trials targeting autophagy are underway to improve ICI-based combination strategies and, consequently, patient outcomes. In particular, randomized phase III trials are ongoing or completed, including ICI doublets such as anti-programmed death-ligand (PD-L1) plus anti-cytotoxic T-lymphocyte-associated protein (CTLA)-4 antibodies (HIMALAYA, NCT03298451) or combinations of ICI with anti-angiogenic therapies such as bevacizumab (IMbrave 150, NCT03434379) or tyrosine kinase inhibitors such as lenvatinib (LEAP-002, NCT03713593) or cabozantinib (COSMIC-312, NCT03755791). Collectively, owing to its unique and original dual mechanism of action, ezurpimtrostat in monotherapy and in combination with ICI constitutes a powerful strategy to improve T cell-mediated immunotherapies. Ezurpimtrostat-based combinations may offer the opportunity to expand the long-term clinical benefits of patients with advanced liver cancer by overcoming the elusive mechanisms of response and resistance to ICI therapy. This strategy is currently being studied in the first-line setting of advanced hepatocellular carcinoma, in the large phase 2 trial ABE-LIVER (NCT05448677), randomizing the combination of the actual standard of care atezolizumab-bevacizumab, with or without ezurpimtrostat.

## Availability of data and material

All data and material avec available

## Competing interests

E.B, M.R, S.M, C.A, E.R and P.H are employees of Genoscience Pharma. E.R and P.H are shareholders of Genoscience Pharma. A.T.R has no conflict of interest to report

## Funding

None

## Authors’ contributions

Conception and design: E. Bestion, M. Rachid, A. Tijeras-Raballand, S. Mezouar

Data analysis: E. Bestion, M. Rachid, S. Mezouar

Study supervision: M. Rachid, E. Bestion, S. Mezouar, P. Halfon

Writing, review of the manuscript: M. Rachid, E. Bestion, G. Roth, T. Decaens, C. Ansaldi, S.

Mezouar, E. Raymond, P. Halfon

## Abbreviations

PPT1: Palmitoyl-protein thioesterase 1
PD-1: programmed death-1
MHC: major histocompatibility complex
HCC: hepatocarcinoma
ICI: immune checkpoint inhibitors
W: Week
CT: Celiac Trunk
HES: hematoxylin-phloxin-saffron
RT: room temperature
PBS: phosphate-buffered saline
PBMC: Peripheral blood mononuclear cell
IFN: interferon
FBS: fetal bovine serum
NBR1: neighbor of *BRCA1* gene 1
TAM: tumor-associated macrophage
PD-L1: anti-programmed death-ligand
CTLA-4: anti-cytotoxic T-lymphocyte-associated protein

## Figure and legends

**Supplementary Fig. 1.**
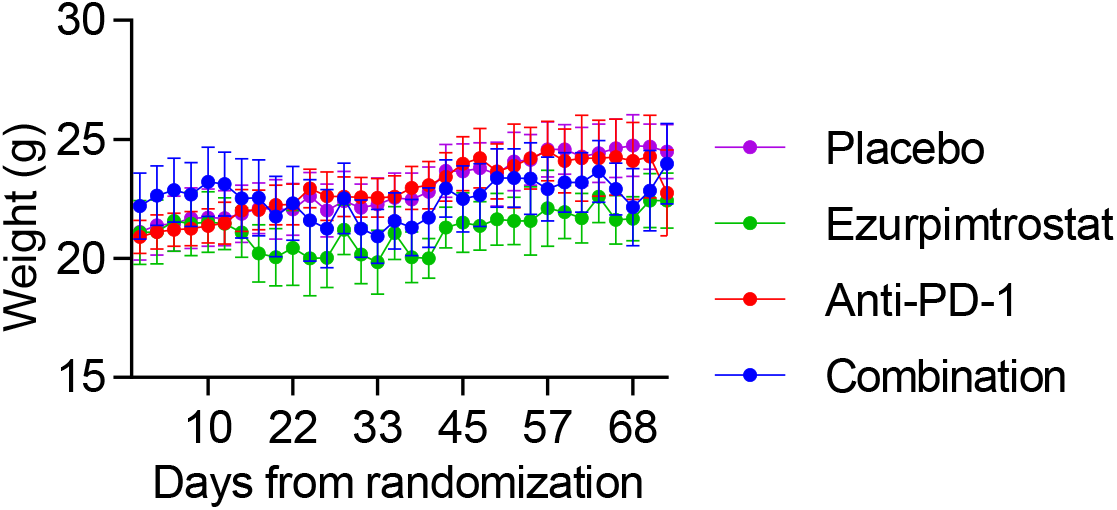
Animal body weight evolution through the study. Representative graph illustrating body weight (g, gram) expressed as the mean ± 95% for vehicle (purple) as control, ezurpimtrostat (green), anti-PD-1 (red), and GNS561 combined with anti-PD-1 (blue).

